# Antithetical contribution of primary and non-primary auditory cortex while listening to speech in noisy scenes

**DOI:** 10.1101/2021.10.26.465858

**Authors:** Lars Hausfeld, Iris M. H. Hamers, Elia Formisano

**Affiliations:** Department of Cognitive Neuroscience, Faculty of Psychology and Neuroscience, Maastricht University, 6200 MD Maastricht, The Netherlands; Maastricht Brain Imaging Centre, 6200 MD Maastricht, The Netherlands; Department of Biomedical Sciences of Cells & Systems, Section Cognitive Neurosciences, University Medical Center Groningen, University of Groningen, Groningen, The Netherlands; Maastricht Centre for Systems Biology, Faculty of Science and Engineering, 6200 MD Maastricht, The Netherlands

**Keywords:** high-field fMRI, audition, auditory scene analysis, speech tracking

## Abstract

Invasive and non-invasive electrophysiological measurements during “cocktail-party”-like listening indicate that neural activity in the human auditory cortex (AC) “tracks” the envelope of relevant speech. Due to the measurements’ limited coverage and/or spatial resolution, however, the distinct contribution of primary and non-primary auditory areas remains unclear. Using 7-Tesla fMRI, here we measured brain responses of participants attending to one speaker, without and with another concurrent speaker. Using voxel-wise modeling, we observed significant speech envelope tracking in bilateral Heschl’s gyrus (HG) and right middle superior temporal sulcus (mSTS), despite the sluggish fMRI responses and slow temporal sampling. Neural activity was either positively (HG) or negatively (mSTS) correlated to the speech envelope. Further analyses comparing the similarity between spatial response patterns in the *concurrent speakers* and *single speaker* conditions indicated that whereas tracking in HG reflected both relevant and (to a lesser extent) non-relevant speech, right mSTS selectively represented the relevant speech signal. Additionally, in right mSTS, the similarity strength correlated with the participant’s comprehension of relevant speech. These results indicate that primary and non-primary AC antithetically process ongoing speech suggesting a push-pull of acoustic and linguistic information.

## Introduction

Speech and other continuous sound streams are increasingly used to examine human auditory processing under naturalistic listening conditions. Using “cocktail-party-like” scenes as stimuli, recent investigations have linked temporally-resolved neural signals, as measured with electrocorticography (ECoG), magnetoencephalography (MEG) and electroencephalography (EEG), to continuously changing features of the input (1). A robust finding is that the envelope of incoming speech is “tracked” by these signals and that, in case of multiple concurrent sounds, the envelope of the relevant (i.e., attended) speech is tracked more reliably compared to the non-relevant one (2–4). These effects have been localized to primary and secondary auditory cortical regions in Heschl’s gyrus and sulcus (HG, HS), superior temporal gyrus (STG), and planum temporale (PT) (2, 5) and, more recently, to subcortical areas (6, 7). However, as these techniques offer limited coverage (ECoG) and/or spatial resolution (EEG/MEG), it has been problematic to distinguish the specific contribution of different auditory brain regions to the neural tracking. The contribution of areas beyond auditory cortex requires further study as well.

To address these issues, here we present ongoing speech stimuli of two speakers (*v1* and *v2*) while measuring brain activity with high-field functional magnetic resonance imaging (fMRI), at high spatial resolution and with whole-cortex coverage. MRI poses challenges for performing auditory studies that increase with field strength mostly due to its noisy and magnetic environment, in particular when presenting long, continuous sound stimuli. Nevertheless, behavioral and EEG speech tracking results from simultaneous MRI and EEG measurements (8) suggested that participants were able to listen selectively to one speaker. Moreover, the EEG-based tracking of the speech envelope inside the MRI scanner was found to be correlated with tracking outside the scanner across participants. It remains unclear, however, whether the hemodynamic signal, an indirect and sluggish measure of neural activity, follows the speech envelope similar to electromagnetic neural signals.

In this study, we investigated fMRI neural tracking of the speech envelope by presenting 5-min blocks of task-relevant speech (Figure 1A) both with and without concurrent speech (referred to as *single speaker* and *auditory scene* condition, respectively; *Method*s *– Sound Stimuli, Experimental Design*). We measured the participants’ (*N* = 15) brain activity with 7T-fMRI (Figure S1; *Methods – Functional MRI*). To avoid potential top-down effects due to repeated sound presentations, each relevant and non-relevant speech segment was presented only once to participants. In addition, to capture the influence of the envelope dynamics, we limited the maximum length of silent periods in the stimuli to 300 ms, which mitigates the strong effect of comparing blood-oxygen level-dependent (BOLD) responses to sound vs. no-sound periods.

**Figure 1.**
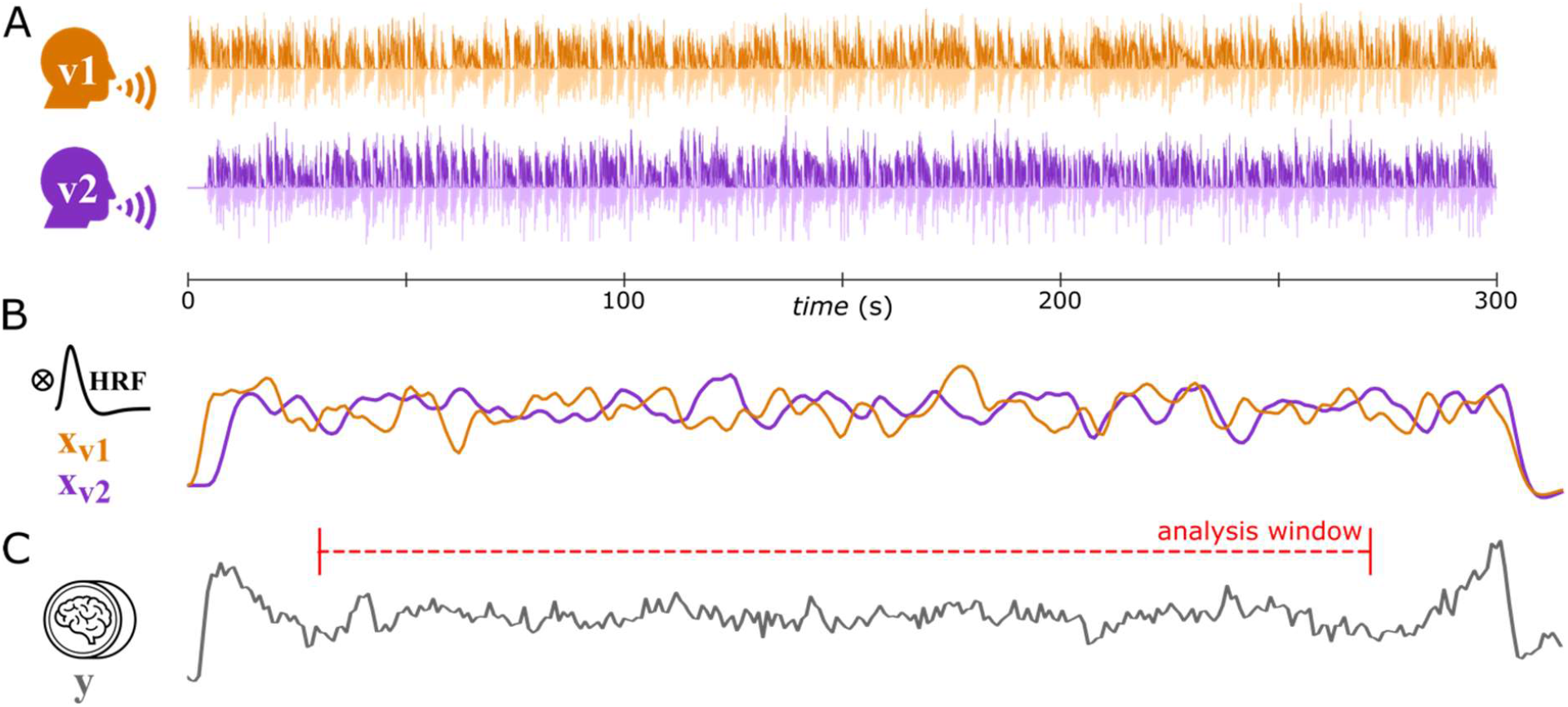
Example Stimuli and Processing. A) Speech waveforms during one trial (300 s) for the auditory scene condition containing speech of speakers v1 and v2. Light shading shows the speech waveform; the saturated lines denote the envelope of these waveforms. Note the initial silent period for speaker v2 introduced to cue and support the tracking of the task-relevant speaker (here: v1). B) To analyze whether the BOLD signal tracked the speech envelope, the envelopes of the relevant and non-relevant speech (see above) were convolved with a hemodynamic response function (HRF, indicated in black). C) BOLD signal for one example region located on HG. Due to the onset and offset effects at the beginning and end of sounds, the speech tracking analysis was limited to 240 s excluding the initial and final 30 s of each block.

Previous tracking studies employed high-temporal precision measurements (EEG, MEG, sampling rate >100 Hz) that allowed analyzing the time series with complex models (fitting >100 parameters). Similar analyses in fMRI are difficult, as the sampling rate is two orders of magnitude lower (TR = 1 s = 1 Hz in this study) and the time-series comparably shorter. To enable speech tracking with fMRI, we acquired long time-courses by presenting listening blocks of 5min (vs. ≤ 60 s in EEG studies) and derived spatial maps of speech tracking within a voxel-by-voxel General Linear Model (GLM)(9) framework. A follow-up multi-voxel pattern analysis (MVPA) of these spatial tracking patterns characterized the relative representation of relevant and non-relevant speech during the auditory scenes

## Results

### Participants follow audiobooks during MRI acquisition

We asked participants to selectively listen to the (relevant) speaker. Participants were asked to answer questions about the audiobook’s content and provide a subjective rating on their selective listening performance after each 5-min segment. The accuracy of responses to content questions indicated that participants were able to listen selectively to the single speaker and auditory scene stimuli (single speaker: 0.760 ± .071 [mean ± s.d.], *t*(14) = 27.736, *p* < .001 [vs. theoretical chance level of 0.25], Cohen’s *d* = 7.16; auditory scene: 0.607 ± 0.175, *t*(14) = 7.877, *p* < .001, *d* = 2.03). This was confirmed by the participants’ subjective ratings (single speaker: 8.72 ± 0.82 [mean ± s.d.]; auditory scene: 5.46 ± 1.67; ratings between 1 and 9: 1 = “could not follow the relevant speaker at all”, 9 = “could follow as well as if presented without noise”). As expected, presenting a second speaker rendered the listening task more difficult (single speaker vs. auditory scene: *t*(14) = 3.672, *p* = .003, *d*_av_ = 1.15), with participants rating their selective listening performance higher during the single speaker vs. auditory scene condition (*t*(14) = 9.208, *p* < .001, *d*_av_ = 2.47).

### Listening to Audiobooks Activates the Speech Comprehension Network

A first analysis showed sustained BOLD signal (de)activation in response to listening blocks compared to baseline, in regions typically involved in speech processing, for both single speaker and auditory scene conditions (Figure S2; *Methods – fMRI Activation Analysis*). Significant activation was observed in auditory cortical regions (HG, STG), superior temporal sulcus (STS) and inferior frontal gyrus (IFG); significant deactivation was found in the temporoparietal junction (TPJ), insula, middle frontal gyrus (MFG) and inferior central sulcus.

The initial investigation of the fMRI data time courses during listening blocks (cf. Figure 1C) revealed early (expected; sound onset) and late (unexpected; preceding sound offset) BOLD signal increases. To remove these tracking-unrelated effects at on- and offset when analyzing the tracking of speech, we restricted our analysis to the central 4-min period of listening blocks by cutting the first and final 30 s (Figure 1C, *Methods – fMRI Tracking Analysis*).

### BOLD responses Reveal Positive and Negative Tracking of Speech

To analyze envelope tracking, we generated predictors for the GLM by convolving the extracted envelopes of the relevant speech and, for auditory scenes, non-relevant speech with a canonical HRF (Figure 1B; *Methods – fMRI Tracking Analysis*). Applying this GLM tracking analysis to the single speaker condition, we found significant fitting and positive parameter estimates (β-values) in bilateral contiguous regions along the HG/HS and on STG both anterior and posterior to HS (Figure 2A). We also found regions with significant fitting and negative β-values in (right) middle superior temporal sulcus (mSTS). These results reveal that low temporal resolution BOLD-fMRI responses track speech amplitude envelopes, extending previous results obtained with high temporal resolution neural measures (EEG, ECoG, MEG).

**Figure 2.**
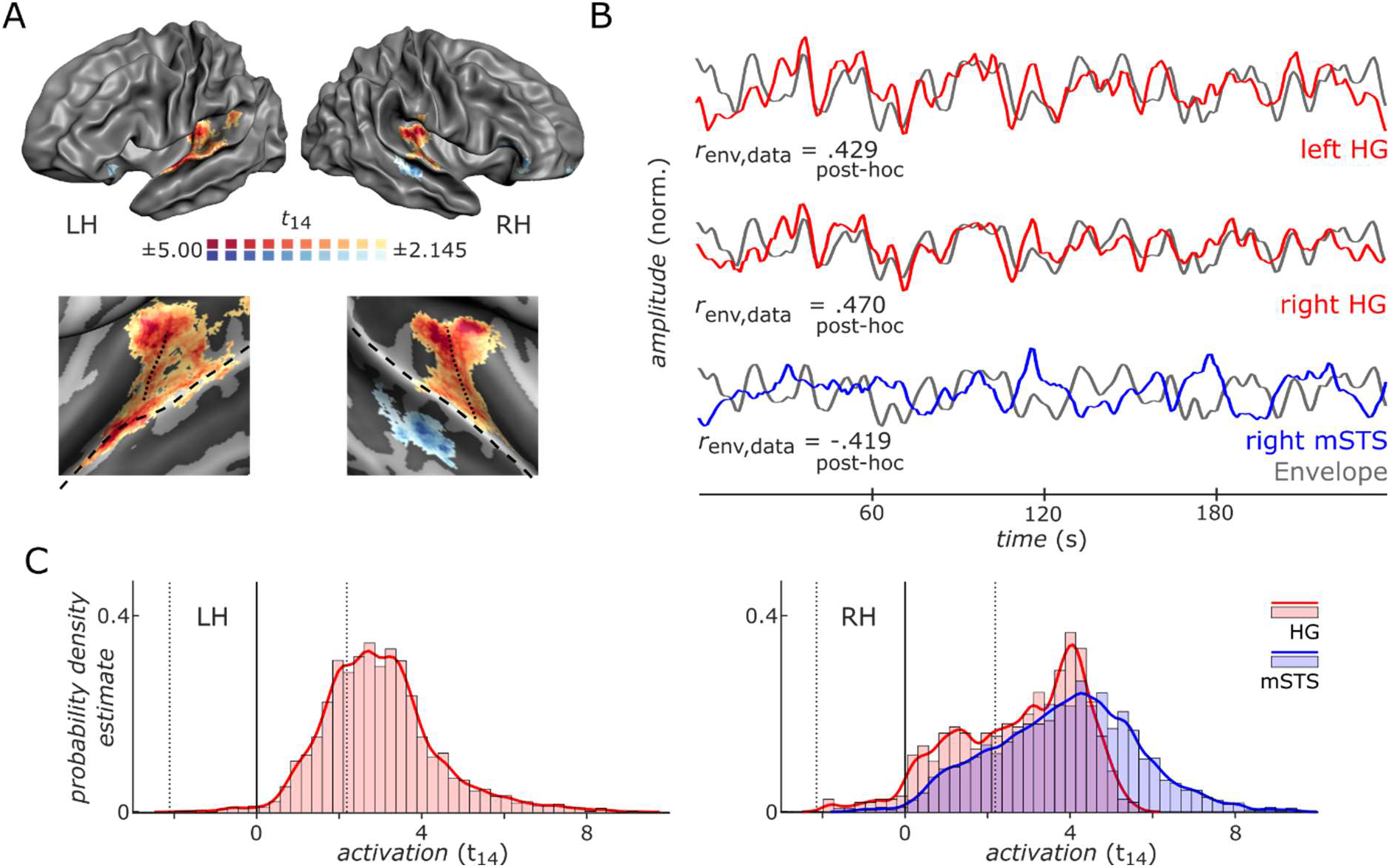
Overview Speech Tracking Single Speaker Condition. A) Color-coding depicts regions showing significant *positive* (warm colors) and *negative* (cold colors) speech tracking in the single speaker condition across participants in the left (LH) and right hemisphere (RH). Upper panels show lateral views of reconstructed average grey-white matter boundary after cortex-based alignment. Lower panels show enlarged views of temporal cortex on the inflated boundary. Dotted and dashed black lines indicate HG and STG, respectively. B) Example BOLD signal time courses (240-s duration, *z*-normalized) averaged across participants during the same block in left and right HG (red line) and right middle STS (blue line) showing significant speech tracking. *Positive tracking* in HG is indicated by the positive temporal correlation between MRI signal and speech envelope (grey lines) whereas *negative tracking* in right mSTS is indicated by their anti-correlation. Significant regions of the group analysis were back-projected to single-participant volume space where the most significant 20% of voxels (non-directional) were selected to create individual time courses. r-values in lower left of each panel indicate the temporal correlation of MRI data and envelope time courses in this example. Note that these correlations are expected given the informed voxel selection and presented here to provide an intuitive interpretation of positive and negative tracking. C) Distribution of statistical values for the “traditional” activation-based analysis of sustained activity in the single-speaker condition as compared to pre-stimulus baseline for the HG and mSTS regions that tracked the speech envelope (see panel A). Significant activation in tracking regions was found in bilateral HG and right mSTS. The distribution of *t*-values is indicated by solid lines and light-colored bars that show the estimate of the probability density function and the normalized histogram, respectively. Statistical maps are thresholded at *p* < .05 (two-tailed) and corrected for multiple comparisons by cluster size (*p* < .05).

The BOLD signal time courses of these regions can be interpreted as showing a positive and negative temporal correlation with the envelope of the speech sound (upper and lower panel of Figure 2B, respectively) which we refer to as *positive* and *negative tracking*. Further analyses showed that areas in bilateral HG displaying positive tracking also displayed significant sustained positive activity with regard to pre-stimulus baseline (i.e., no sound)(Figure 2C, red color). Interestingly, we also found positive sustained activity for areas in right mSTS that displayed negative tracking, (Figure 2C, blue color). This indicates that a positive BOLD response to sound with respect to pre-stimulus baseline can show both positive and negative tracking of the speech envelope. To verify that the negative correlation in mSTS is not due to the specific choice of hemodynamic response model, we varied the HRF model for a wide range of values of the time-to-peak parameter (3.5-7 s) (Figure 3A). Results showed that HRF models with shorter time-to-peak fitted better in medial HG, whereas HRF models with longer time-to-peak fitted better in mSTS (Figure 3B). Importantly, the tracking of the envelope in mSTS remained negative for the entire range of HRF models considered (Figure 3B). The results presented here were obtained with a time-to-peak parameter of 4.5 s, providing a compromise between fitting BOLD responses in both the HG and mSTS regions.

**Figure 3.**
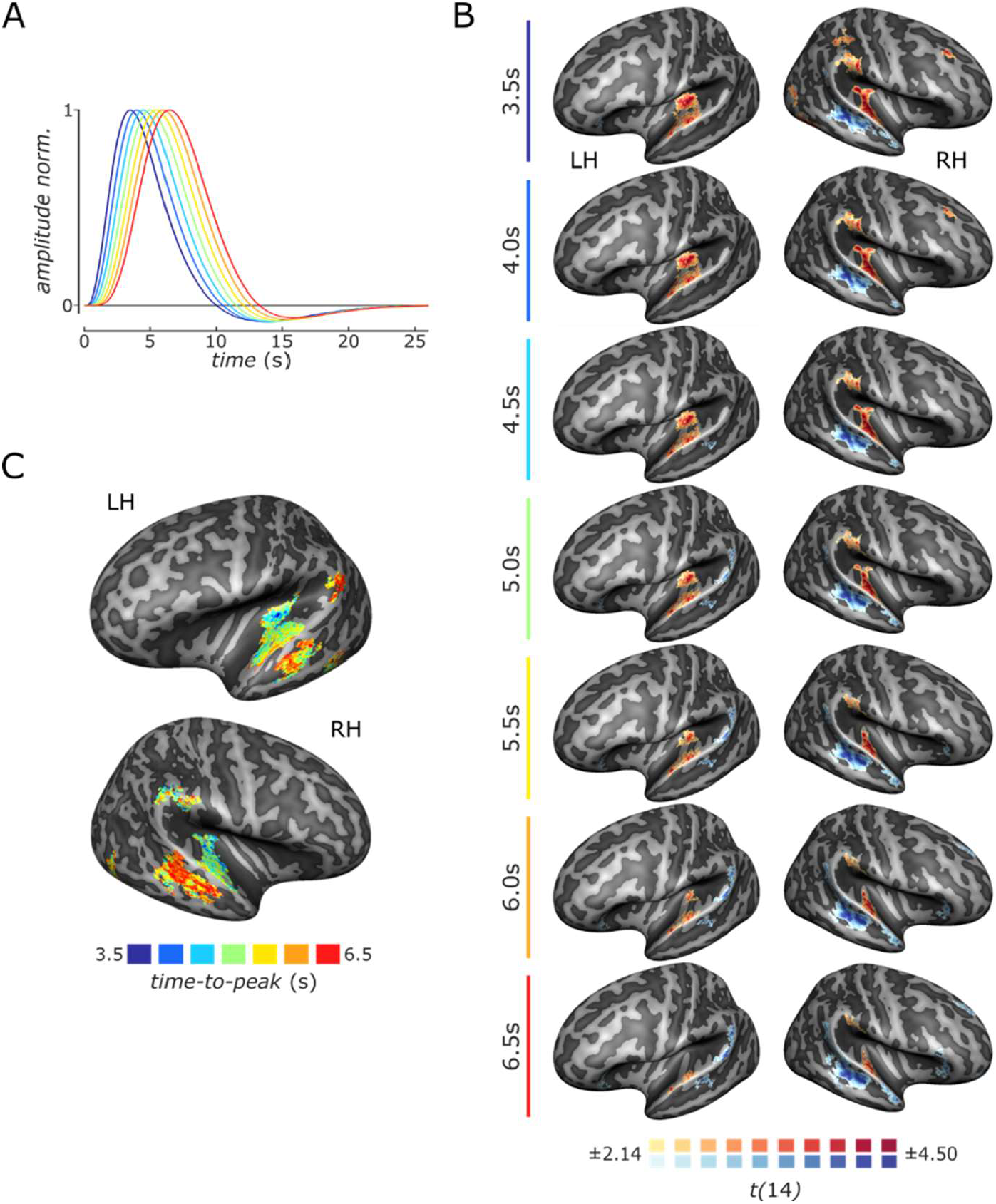
Time-to-Peak Dependency of Speech Tracking. A) Curves present 7 hemodynamic response functions (HRFs) created by varying the time-to-peak latency, a crucial parameter when specifying the HRF for analysis via GLM. Colors indicate the HRFs with different time-to-peak parameters (see panel C) for color coding). B) Results of speech tracking for the single speaker condition for analyses with the different HRFs. Color-coded regions show significant speech tracking across participants (*p* < .05, multiple comparison corrected by cluster size). Maps are presented on inflated reconstructions of the average grey-white matter boundary after cortex-based alignment. C) Overview of the time-to-peak parameter resulting in best speech tracking performance. These maps (upper panel: left hemisphere; lower panel: right hemisphere) are restricted to loci showing significant speech tracking for at least one model.

### Tracking Patterns Reveal (Non-)Relevant Speech Processing in HG and mSTS Regions

Previous research highlighted that spatial activation patterns across the (auditory) cortical surface represent auditory objects including speech streams and that these spatiotemporal representations are significantly affected by selective attention in multi-talker scenes (3, 5, 10–13). We thus performed a spatial pattern similarity analysis to investigate the effects of selective attention in the regions-of-interest identified during the single speaker condition (i.e., bilateral HG and right mSTS regions). Specifically, we compared spatial maps for tracking (i.e., voxel-wise parameter estimates for speech envelope predictors) obtained from the single-speaker condition (Figure 2A) with those from the auditory scene condition, obtained by including amplitude envelope predictors of the non-relevant speech in addition to the relevant speech (Methods – Spatial Pattern Analysis of Tracking Maps).

Overall, our multivariate results show that the pattern similarity of tracking maps across tasks was significant in the left and right HG regions for relevant speech (Figure 4A, red; *p*_adj_ < .001; *t*(14) > 4.665, Cohen’s *d* > 1.20; multiple comparison corrected [MCC] across 6 tests vie False-Discovery-Rate [FDR](14)) and for non-relevant speech in right but not left HG (right: *p*_adj_ = .017; *t*(14) = 2.567, *d* = 0.66; left: *t*(14) = 1.094, *p*_unc_ = .146). In the right mSTS region, we found significant similarity of tracking patterns for relevant speech (Figure 4A, blue; *t*(14) = 3.24, *p*_*adj*_ = .006, *d* = 0.84) but not for non-relevant speech (*t*(14) = -0.295, *p*_unc_ = .614, *d* = 0.10). This indicates similar BOLD speech tracking maps when listening to a single speaker or an auditory scene of two concurrent speakers for relevant speech in the HG and right mSTS regions and for non-relevant right HG.

**Figure 4.**
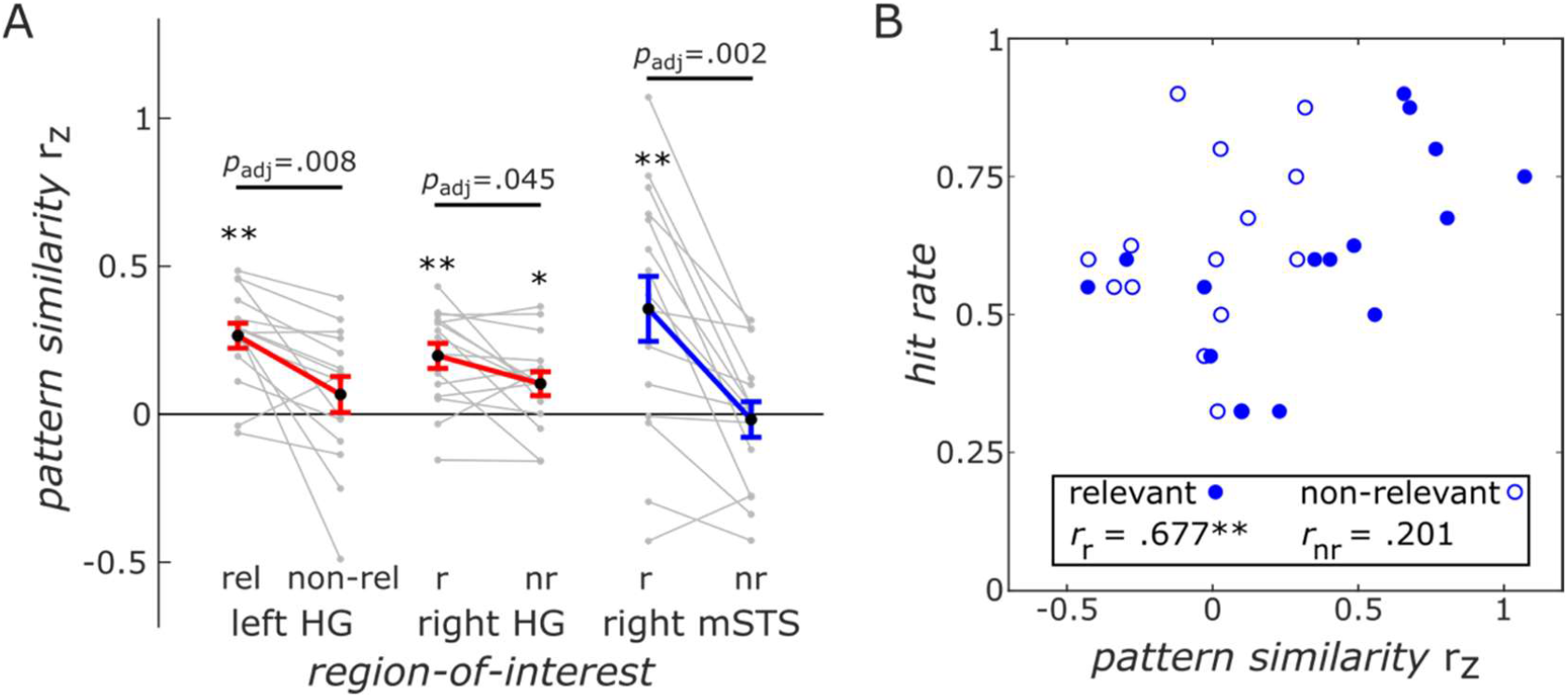
Analysis of Pattern Similarity between Tracking Maps of Single Speaker and Auditory Scene Conditions in Regions-of-Interest. A) Similarity of spatial patterns of speech tracking between the single speaker and auditory scene condition for the HG (red) and mSTS (blue) regions determined by significant tracking for the single speaker conditions (see Fig. 1A). Grey lines indicate single participants and thick coloured lines their average (error bar ± s.e.m.). Asterisks denote significant pattern similarity between single speaker and scene conditions (\*\**p*_*adj*_ < .01, *\*p*_*adj*_ = .017, two-tailed; FDR-adjusted p-values; MCC across 6 tests) and straight lines show differences in pattern similarity (MCC across 3 tests) between the relevant speech (r/rel) and the non-relevant speech (nr/non-rel). B) Scatter plots showing pattern similarity in the right mSTS region and behavioural performance. Circles represent data points of individual participants for relevant (filled) and non-relevant speech (open). A non-parametric correlation analysis across participants (\*\**p*_adj_ < .05, two-tailed; MCC across 6 tests) supported a positive relationship between pattern similarity for tracking patterns for relevant speech and participants’ responses. We did not find a relationship for non-relevant tracking patterns. Similar analyses in HG (not shown) did not show significant associations.

### Tracking Patterns Reveal Dominant Processing of Relevant Speech

In the left HG region, the similarity of tracking patterns was modulated by relevance (Δ*r* = 0.199; *t*(14) = 3.297, *p*_adj_ = .008, *d*_av_ = 1.00; MCC across 3 tests). Moreover, when analyzing the tracking map similarity in the right hemisphere regions, we found that the tracking map similarity was affected by relevance (*F*(1,14) = 28.45, *p* < .001, 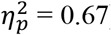) but not region (*F*(1,14) = 0.064, *p* = .804, 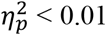). However, these factors interacted significantly (*F*(1,14) = 8.079, *p* = .013, 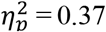; repeated-measures ANOVA). Post-hoc analyses showed that the pattern similarity between tracking maps in the right mSTS region of the single speaker condition was more similar to tracking maps for relevant speech compared to non-relevant speech (*p*_adj_ = .002; *t*(14) = 4.510, *d*_av_ = 1.14). Note that right HG showed a weaker (HG: Δ*r* = 0.094; mSTS: Δ*r* = 0.374) but similar modulation by relevance for the similarity of tracking maps (*p*_adj_ = .045; *t*(14) = 2.203, *d*_av_ = 0.59).

### Tracking Patterns Link to Relevant Speech Comprehension in mSTS but not HG regions

To examine whether the similarities of spatial tracking patterns were linked to behavior, we performed a correlation analysis between fMRI (i.e., tracking patterns for relevant and non-relevant speech) and behavioral data (accuracy of answers to content questions). Across participants, both left and right HG regions did not show significant correlation (left HG: *r*_s_ = -.134, *p*_unc_ = .625; right HG: *r*_s_ = .004, *p*_unc_ = .987; Spearman’s correlation coefficient *r*_s_, two-tailed, significance assessed by permutation test, *n*_perm_ = 10^5^). In contrast, the negative tracking region in right mSTS showed a significant brain-behavior relationship (Figure 4B). More specifically, the tracking pattern similarity for single speech and relevant speech was positively correlated with the response accuracy to content questions (*r*_s_ = .677, *p*_unc_ = .007; *p*_*adj*_ = .041, MCC-adjusted across 6 tests) while this did not hold for the tracking pattern similarity for non-relevant speech (*r*_s_ = .201, *p*_unc_ = .466). The brain-behavior correlation in mSTS was significantly stronger for pattern similarities for relevant vs. non-relevant speech (*p* = .032, two-tailed, permutation test, *n*_perm_ = 10^5^).

Overall, these results show that speech relevance modulated the similarity of tracking maps in HG and mSTS regions from the auditory scene to the single speaker condition with higher similarity for the relevant vs. non-relevant speech. Furthermore, our observations indicate that right mSTS reflects relevant but not non-relevant speech and that this similarity of tracking maps for relevant speech was linked to its comprehension.

## Discussion

In this study, we measured cortical responses to continuous speech stimuli using high-field fMRI. In a first step, we showed that the hemodynamic response follows or “tracks” the speech envelope amplitude. More specifically, we found that the BOLD signal tracks the ongoing speech envelope of a single speaker in bilateral HG, STG and STS. These findings resemble the speech tracking observed by direct and temporally resolved neural measures (ECoG, MEG and EEG), which showed robust tracking of the speech envelope amplitude.

Interestingly, our results showed positive and negative tracking of the speech envelope by the HG and mSTS regions, respectively. Both positive and negative tracking was found in areas that showed increased sustained activity in response to speech sounds (i.e., the speech envelope modulated the signal “on the plateau” of the positive BOLD activation). We interpret the positive tracking in the HG region to reflect the ongoing envelope amplitude of the speech stream. This is in line with previous ECoG studies showing that activity in HG and middle STG is correlated with responses to speech (3, 5, 15). We interpret the negative tracking observed in the right mSTS region to reflect cortico-cortical top-down signals that aid in following relevant speech in particular during periods of low speech audibility, i.e., when the task is more difficult (e.g., due to the fMRI noise and lower intensity of relevant speech).

Layer-resolved high-field MRI acquisitions might help to better define the role of this region in terms of top-down and bottom-up input when listening to a single speaker or an auditory scene (e.g., De Martino et al., 2018). Whether these regions of BOLD speech tracking coincide with the sources of the speech tracking observed with neuro-electromagnetic signals remains an open question. The relationship between neural activity, electric and hemodynamic signals is complex (17–19). Concurrent measures of speech tracking by EEG and fMRI (8) would allow linking, within participants, the observed results by hemodynamic and neuroelectric measures more directly and shed light on the underlying neural processes.

After having established that hemodynamic signals follow continuous speech signals, we performed an analysis of multi-voxel patterns that revealed high similarity between spatial tracking patterns of relevant speech in an auditory scene and speech presented without concurrent distractor in HG and the STS regions. In right mSTS, this similarity of the tracking patterns was (positively) correlated with the assessment of speech comprehension across participants.

Examining the spatial patterns of speech tracking maps for the HG region, we found a high similarity between the single speaker and auditory scene conditions for both the relevant and – to a lesser extent – non-relevant speech. These results indicate that the overall incoming speech signal, containing the relevant and non-relevant speech, is reflected in the HG region including medial and lateral HG/HS and adjacent STG. In these regions, the pattern similarity was marginally higher for relevant speech in comparison to non-relevant speech (Figure 4A). This is in line with previous observations showing attentional modulation of envelope representations with high selectivity for attended vs unattended speech using ECoG (3, 5) and MEG (20, 21) in these regions. Although being significantly lower compared to relevant speech, the pattern similarity of tracking maps for non-relevant speech suggests residual information about non-relevant speech in the HG region.

In contrast, for the right mSTS region, we found significantly higher pattern similarity of tracking maps of relevant speech vs. non-relevant speech tracking and no significant similarity for tracking maps of non-relevant speech. In addition, we observed that participants with a higher pattern similarity between single speaker and relevant speech of the auditory scene performed better in a comprehension task about the audiobook’s content. This result might indicate that right mSTS processes exclusively information reflecting the relevant speech, implying that the cocktail party is resolved at this stage. This fits with previous results in which pattern similarity in mSTS/STG only represents relevant speech but not non-relevant speech or music; the effect in this region was category specific such that relevant speech but not music showed significant pattern similarity (11). Another possible explanation for these results is that activation in this area reflects increased top-down control of selective listening at a temporal scale of envelope changes (i.e., phonemes, syllables and words; (22, 23)) with decreasing amplitude of the relevant speaker and thus increasing energetic masking. These explanations are not mutually exclusive such that both bottom-up and top-down contributions important for selective listening are represented in this region’s signals. This region partially overlaps with electrophysiological recording sites in STG, which suggested responses to sustained features of the speech signal (i.e., speech envelope)(15) in line with the current findings. Results of a recent fMRI study using continuous speech stimuli suggested that the HG region mostly represented spectral information (related to the envelope amplitude), while the mSTS region was mostly correlated with semantic features (24). While this is in agreement with our findings in the HG region, the significant tracking of the envelope in mSTS presumably tracking phonological or semantic features might be explained by correlations between these features and the amplitude envelope. Additional differences, for example in analyses or data acquisition (3T vs. 7T MRI, 1Hz vs 0.5Hz sampling) could explain these observations. However, overall, the current and previous results support that the mSTS region links bottom-up acoustic and top-down linguistic processing of relevant speech during auditory scenes.

The correlation between tracking map similiarites in mSTS and participants’ performance on answering content questions suggests that activity in this region reflects the behavioral outcome. It might be linked to previous observations associating MEG and EEG-based speech tracking with speech intelligibility and comprehension (4, 25–27) although, if analyzed, this brain-behavior association is not always found (28). The STS has been linked to intermediate linguistic representations (23) and EEG-based speech tracking exploiting linguistic features was correlated with the performance in speech comprehension tasks across participants (29). Our findings corroborate these observations by suggesting a neural source of this link between behavior and linguistically informed speech tracking.

Our analyses showed similar tracking patterns for non-relevant speech in the HG regions. Previous studies using EEG, MEG and ECoG have indicated that background sounds including speech are represented in the auditory system in particular at early latencies of processing (30–33), which is in line with the current finding suggesting information of non-relevant speech being represented in earlier areas in auditory cortex and being represented less in higher areas in the auditory processing hierarchy like STG and STS.

To conclude, our results showed that speech tracking, a robust phenomenon observed with high temporal resolution and neuro-electric signals, can be observed with low temporal sampling and high-field fMRI BOLD responses. Furthermore, we found opposite tracking of speech in HG and mSTS regions and tracking of non-relevant speech in HG but not mSTS. Speech tracking in the mSTS was linked to speech comprehension. These results indicate neural processes potentially related to stronger feedback and linguistic integration processing in mSTS compared to HG aiding successful listening in noise. In addition, these results provide support for neural signals in mSTS that reflect a processing stage at which the cocktail-party is resolved.

## Acknowledgements

We would like to thank Federico De Martino for help with data acquisition and comments on the manuscript, Anne Bach for help with participant recruitment, and our participants for their collaboration and time. This work was supported by Maastricht University, the Dutch Province of Limburg (E.F.), and the Netherlands Organization for Scientific Research (NWO; VENI grant 451-17-033 to L.H.).

## Materials and Methods

### Participants

Fifteen students (native German speakers) of Maastricht University (13 female, 2 male, mean age ± [s.d.]: 24.1 ± [3.8] years, age range: [19 33] years), after signing the written informed consent, took part in the experiment and received course credit or gift vouchers for their participation. The local ethics committee of the Faculty of Psychology and Neuroscience (*Ethics Review Committee Psychology and Neuroscience*) at Maastricht University approved the experimental procedures of the study (#167_09_05_2016).

### Sound Stimuli

We presented participants with speech (audiobook excerpts)(31) of one female (*v1*; *f*_*0*_ = 159 ± 8.3 Hz, mean ± s.d.) and one male speaker (*v2*; *f*_*0*_ = 107 ± 7.3 Hz). The fundamental frequency *f*_*0*_ for each excerpt was determined by averaging *f*_*0*_ contours obtained with the YIN algorithm (34). Sounds were played on top of MRI scanner noise and delivered via an MR-compatible sound system (Sensimetrics S14, Sensimetrics Corporation, Malden, MA) diotically by in-ear earphones. Sound stimuli were presented at a high but comfortable level that was individually adjusted at the beginning of the experiment. Sound intensity of the two audiobooks was equalized based on root-mean-square (RMS), i.e. v1-speech was presented at a signal-to-noise ratio (SNR) of 0 dB_RMS_ with regard to v2-speech. To avoid clicks, the onset and offset of each speech signal were ramped (linear ramps of 0.1 s). Auditory stimuli were digitized using a sampling rate of 44.1 kHz and 16 bits. For all sound stimuli, silent periods (e.g., during words or sentences) were adjusted to a duration of at most 300 ms using Praat (35).

### Experimental Design

The design included two conditions, 1) the *single speaker* condition, i.e. the presentation of speech of one audiobook, and 2), and the *auditory scene* condition, i.e. the concurrent presentation of speech of two audiobooks. To obtain sufficient samples for each presentation, each block lasted 5 min. During the auditory scene condition, the target speech started 4.5 s before the distractor speech to provide listeners with an auditory cue indicating the target speech (see Figure 1).

### Functional MRI

Brain imaging was performed with a 7-Tesla Siemens Siemens Magnetom scanner with a whole brain coil at the Maastricht Brain Imaging Center (Maastricht, The Netherlands). Anatomical scans were acquired during each session with an MP2RAGE sequence (36)(voxel size: 0.65mm isotropic; 240 slices; FoV: 208 mm; TR: 5000 ms; TE: 2.51 ms; GRAPPA 2) and masked with the second inversion contrast. For each participant, 6 functional runs of 722 ± 10 volumes (mean ± s.d.; range [698 756]) with whole cortex coverage were collected using an echo-planar imaging (EPI) sequence with multiband 3 acceleration (57 slices; voxel size: 1.5 mm isotropic; FoV = 192 x 192 mm; TR = 1000 ms; TE = 19 ms; GRAPPA 2). For correcting EPI distortions two sets of five images were acquired in opposite phase encoding directions (i.e., anterior-posterior and posterior-anterior) between the third and fourth functional runs.

The two conditions (i.e., single speaker and auditory scene) were presented in different runs (single speaker in runs 1 and 4, auditory scene in runs 2, 3, 5 and 6) each containing one block of the v1-task and v2-task with alternating first condition counter-balanced across participants (Figure S1). Participants were asked to selectively listen to speech of v1 (*v1-task*) or v2 (*v2-task*). Presentations for the two conditions included a 15-s rest period followed by the 5-min presentation of the sound stimulus and was followed by another rest period of 10 s (a fixation cross was presented throughout in the center of the visual display through a mirror at the back of the scanner). Subsequently, participants indicated their subjective task performance (“How well did you follow the relevant voice?”, range: 1 [could not follow the relevant speaker at all] – 9 [could follow as well as if presented without noise]) and responded to five questions on the content of the (relevant) audiobook (4-alternative-forced-choice task; answer alternatives indicated by A, B, C, D) by button press (31).

### Data preprocessing

Preprocessing of both functional and anatomical data was performed with BrainVoyager (v21.4, Brain Innovation, Maastricht, The Netherlands). FMRI data preprocessing consisted of slice-scan-time correction, motion correction, EPI distortion correction, and temporal high-pass filtering (11 cycles per run ≈ 0.015 Hz). EPI distortions were corrected using BrainVoyager’s COPE plugin (37)(v1.1). Functional runs were individually aligned to anatomical scans and transformed to Talairach space (38). The functional data was spatially smoothed (4 mm FWHM) and individual maps were projected onto the group-aligned surface via cortex-based alignment (39) to create group maps.

### fMRI Activation Analysis

To detect cortical regions responding to the presentation of and selectively listening to audiobooks in comparison to pre-stimulus baseline, the functional data of the single speaker and auditory scene conditions was analyzed using a general linear model (GLM). Because we observed strong onset and offset effects and were interested in the sustained activation for the tracking analysis (see fMRI Tracking Analysis), listening blocks were modelled by three predictors reflecting the onset, sustained and offset responses.

### fMRI Tracking Analysis

Before analysis, the functional data was cut to 4min per block by removing the initial and final 30s to avoid confounds from onset or offset effects observed during data exploration (Figure 1C). Subsequently, the functional data was analyzed - voxel-by-voxel - for the tracking of speech envelopes by making use of the GLM framework. More specifically, we modelled BOLD voxel time courses by **y** = **Xβ** + **ε** (where y denotes a voxel time course, X a design matrix, β coefficients and ε the error term). The design matrix included the main predictors for the speech envelope time courses and confound predictors reflecting participant’s motion, level difference between first and second block within each run, and an offset (constant). For the single speaker condition, one predictor reflecting the presented speech was included. For the auditory scene condition, two predictors were included, one for relevant speech and one for non-relevant speech.

### Spatial Pattern Analysis of Tracking Maps

To investigate BOLD activity during listening to auditory scenes, we analyzed the spatial patterns. For this pattern similarity analysis, the tracking maps in the HG and mSTS regions from the analysis of single speaker tracking were used as templates. The pattern similarity was computed between the tracking maps in the same regions obtained from BOLD activity during auditory scene presentations for each relevant voice using Pearson’s correlation and Fisher’s transformation 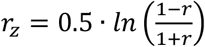. Statistical testing of these scores within and across regions was done via repeated-measure ANOVAs and paired *t*-tests. For ANOVAs and paired *t*-tests, effect sizes were estimated using partial η square 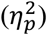 and Cohen’s *d* using the averaged variance in the denominator 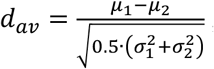, respectively.

## Supplementary Figures

**Figure S1.**
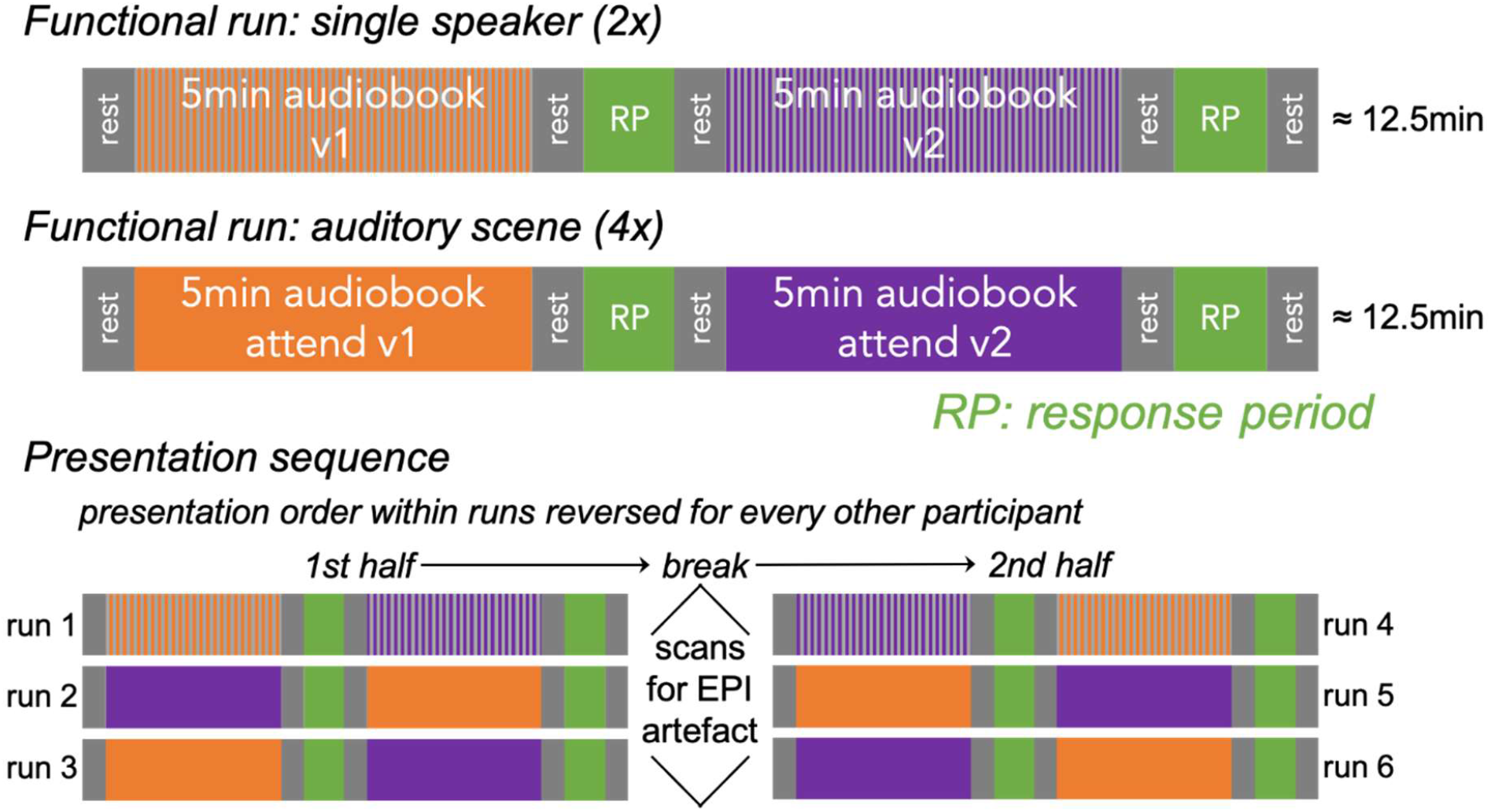
Overview of MRI-Design. The upper plot shows examples of functional runs for the single speaker (repeated twice during each scanning sessions) and auditory scene conditions (repeated four times). After presentation of the single speaker or auditory scene stimuli (5 min) and a rest period (15 s), participants were asked to answer 5 questions on the presented content and to rate their listening performance. After another rest period, another single speaker or auditory scene conditions was presented followed by the response period. The lower plot shows the overview one scanning session. These were split in two halves by scans used for EPI artefact correction after 3 functional runs.

**Figure S2.**
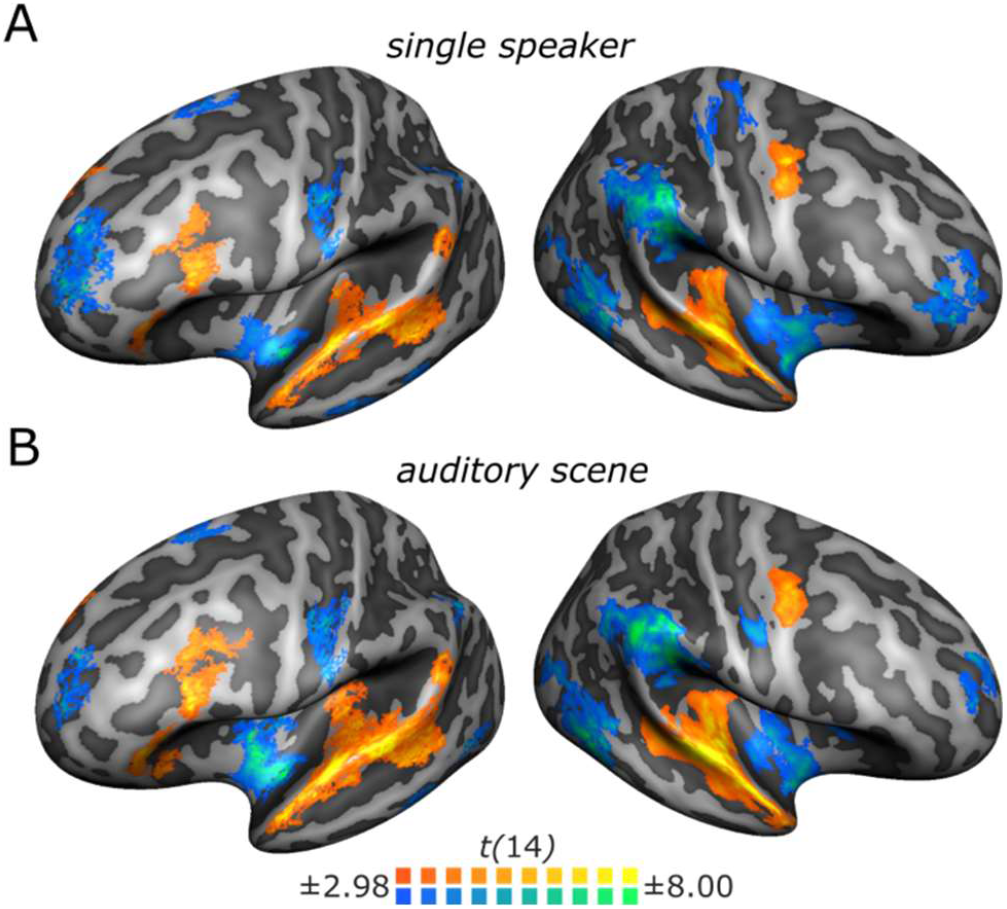
Sustained Activation and Deactivation during Listening Blocks. A) Cortical Maps show regions with significant sustained responses during listening blocks as compared to baseline periods (i.e., no sound) during the single speaker condition for the right and left hemisphere. B) Same as A) but for the auditory scene condition. Statistical maps are thresholded at *p* < .01 (two-tailed) and corrected for multiple comparisons by cluster-size. GLMs included, in addition, predictors coding for onset and offset responses (maps not shown).

